# On the limit: Working memory capacity constrains learning of complex visuomotor mappings

**DOI:** 10.64898/2026.01.30.702906

**Authors:** Vikranth R. Bejjanki, Jordan A. Taylor

## Abstract

Human sensorimotor adaptation critically depends on the ability to map sensory inputs onto motor outputs. While this process was once thought to rely primarily on cerebellar-dependent implicit recalibration, recent work has revealed that explicit, cognitive strategies, relying on working memory and executive function, play a substantial role. However, it remains unclear whether explicit strategies can scale to support learning of complex sensorimotor mappings. Here, we parametrically tested the capacity of explicit strategies to solve a complex visuomotor rotation task by varying the number of target–rotation pairings that participants had to acquire. In experiment 1, participants were tasked with learning four visuomotor mappings – well within working memory capacity, based on prior studies. We found that participants achieved near-perfect compensation, which was best explained by the retrieval of stored target–rotation associations, rather than by computationally demanding algorithmic strategies. In experiment 2, in an attempt to push beyond working memory capacity, participants were tasked with learning eight mappings. Unlike in experiment 1, here we found that participants failed to fully compensate for the rotations, reaching asymptotic performance of only ∼50%, despite evidence of continued strategic engagement. This performance limit was fully predicted by a parameter-free working-memory model where performance is a mixture of a fixed number of stored target-rotation associations and random guessing. These findings reveal a cognitive “bandwidth limit” on the effectiveness of strategies for sensorimotor adaptation: when task complexity exceeds this limit, adaptation plateaus, defining a fundamental constraint on how far higher-order cognition can go to support learning.

**Significance Statement:** Human learners often compensate for perturbations in sensorimotor mappings by deploying explicit, cognitive strategies, but the scalability of these strategies for more complex tasks remains unknown. Here, we tested how performance changes as people must simultaneously learn an increasing number of distinct visuomotor rotation mappings. When presented with only a small set, participants have little difficulty – they learn by opting to forgo computationally demanding algorithmic strategies in favor of storing and retrieving target-specific solutions from working memory. However, as the number of mappings grows, this response caching strategy fails, with performance plateauing in a manner that is consistent with a capacity-limited memory bottleneck. These results highlight the critical constraint that working memory imposes on the utility of explicit strategies for sensorimotor adaptation in complex tasks.

## Introduction

Humans are capable of not only learning sophisticated and even arbitrary sensorimotor skills, but also flexibly adapting these skills to changing conditions. This ability requires learners to discover and adjust complex action-outcome mappings, so that motor commands continue to be effective even in unfamiliar settings. While this process of sensorimotor adaptation has traditionally been attributed to a unitary implicit process (Jordan & Rumelhart, 1992; Shadmehr et al., 2010; Wolpert & Ghahramani, 2000), it has become increasingly clear that much of the observed performance improvements stem from deliberate, explicit strategies (Huberdeau et al., 2015; McDougle et al., 2016; Tsay et al., 2024).

For instance, studies using visuomotor rotation tasks – where visual feedback is rotated relative to hand movement – have shown that implicit recalibration appears to be highly inflexible: it is largely insensitive to perturbation magnitude, saturates below full adaptation, is context independent, and is incapable of consolidating multiple sensorimotor mappings (Bond & Taylor, 2015; Kim et al., 2018; Morehead et al., 2017; Schween et al., 2019; Schween et al., 2018; Wilterson & Taylor, 2021). In contrast, explicit strategies have been shown to be remarkably flexible to task demands, are seemingly essential for rapidly improving performance, and may even account for the lion’s share of adaptation (Bond & Taylor, 2015, 2017; Fernandez-Ruiz et al., 2011; Haith et al., 2015; McDougle et al., 2016; McDougle & Taylor, 2019; Schween et al., 2019; Schween et al., 2018; Wilterson & Taylor, 2021). However, using an explicit strategy to overcome even simple perturbations places significant demands on working memory and executive function (Anguera et al., 2010; Areshenkoff et al., 2024; Christou et al., 2016; Galea et al., 2010; McDougle & Taylor, 2019; Taylor & Ivry, 2014; Velazquez-Vargas & Taylor, 2024). This raises the question: to what extent can explicit strategies handle increasingly complex sensorimotor perturbations?

The extent to which explicit strategies can support adaptation likely depends on their underlying form – explicit strategies can be broadly categorized into algorithmic and retrieval-based modes of operation (Georgopoulos et al., 1989; Georgopoulos & Massey, 1987; Georgopoulos & Pellizzer, 1995; Logan, 1988; McDougle & Taylor, 2019; Pellizzer & Georgopoulos, 1993). Algorithmic strategies, such as visuomotor mental rotation, involve mentally simulating potential aiming solutions to counteract the perturbation. However, they come with a cost: preparation time linearly increases with rotation magnitude (Bhat & Sanes, 1998; McDougle & Taylor, 2019; Shepard & Metzler, 1971; Stransky et al., 2010), indicating higher computational demands. Alternatively, participants could avoid this computational cost by retrieving a previously successful solution from working memory (Anguera et al., 2010; Fresco et al., 2023; Haith & Krakauer, 2018; Velazquez-Vargas & Taylor, 2024). However, retrieval strategies appear to be subject to significant working memory capacity constraints: participants may be able to cache strategies when task complexity is low, but this may break down as complexity increases (Velazquez-Vargas & Taylor, 2024). These working memory limits for explicit strategies may stem from capacity constraints of visual short-term working memory (Cowan, 2001, 2010; Luck & Vogel, 1997; Zhang & Luck, 2008), somatosensory memory (Ebrahimi & Ostry, 2024; Sidarta et al., 2018), or a distinct form of motor working memory (Hillman et al., 2024; Hillman et al., 2025; McDougle & Hillman, 2025).

Such constraints call into question the true usefulness of explicit strategies for sensorimotor adaptation beyond relatively simple learning situations. For instance, successful adaptation has only been observed in studies demanding one or a few strategies (McDougle & Taylor, 2019; Velazquez-Vargas & Taylor, 2024). In contrast, even the most common sensorimotor mappings, from simply controlling a limb to playing a musical instrument, are much more complex. Little research has examined how variables such as task complexity influence such learning – i.e., while it is clear that explicit retrieval strategies are critically constrained by complexity, due to their dependence on executive function and working memory capacity, no prior study has parametrically examined the manner in which these constraints play out in visuomotor tasks as a function of complexity.

Here, we examine the ability of explicit strategies to handle a complex learning task – one that goes beyond previous studies and would be a step closer to the challenges of everyday motor behavior. Specifically, participants were tasked with adapting to a complex set of perturbations through explicit strategies alone. We show that as the complexity of the mapping increased, explicit adaptation broke down. This failure did not stem from limitations in participants’ ability to deploy algorithmic strategies; instead, participants’ behavior was consistent with an inability to appropriately cache the complex set of action-outcome mappings in working memory. These findings highlight how working memory capacity might represent an upper bound on the utility of explicit strategies for sensorimotor adaptation.

## Results

While explicit strategies have been shown to play a major role in sensorimotor adaptation, their dependence on capacity-limited cognitive resources could limit how well they can handle complex mappings. To address this, we tasked participants with learning to counteract a complex set of perturbations in a visuomotor rotation task, where each target location was associated with a unique visuomotor rotation (Fig. 1). This is a variation of the classic dual-adaptation paradigm which, here, requires either learning four (Experiment 1) or eight (Experiment 2) pseudo-randomly interleaved, spatially-dependent action-outcome mappings. Importantly, we delayed cursor feedback in each trial to suppress implicit recalibration processes, so that learning of the complex sensorimotor mapping would be the result of explicit aiming strategies.

**Figure 1.**
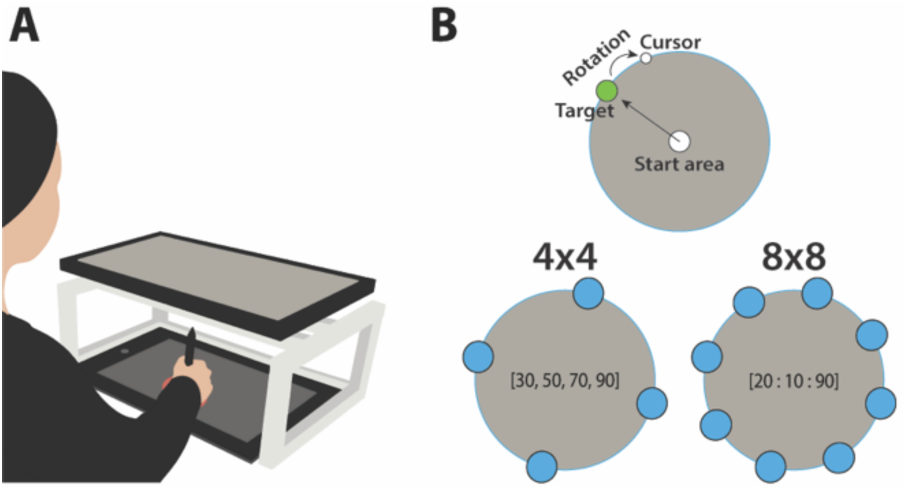
Visuomotor Rotation Task. (A) Experimental Apparatus. Participants were tasked with bringing a virtual cursor to a target by sliding a stylus on a digital tablet positioned under a monitor, which occluded vision of the limb. (B) *Top*: Schematic showing a typical trial. *Bottom:* Target locations and rotation magnitudes in the two experiments. In experiment 1, participants reached to one of four targets (located at 15°, 105°, 195° or 285°), each randomly associated with a different visuomotor rotation magnitude (±30°, ±50°, ±70° or ±90°). In experiment 2, participants reached to one of eight targets (located at 15°, 60°, 105°, 150°, 195°, 240°, 285° or 330°), each randomly associated with a different visuomotor rotation magnitude (±20°, ±30°, ±40°, ±50°, ±60°, ±70°, ±80°, or ±90°).

### Experiment 1

To familiarize participants with the general task, participants were initially presented with a Baseline block where they experienced veridical cursor feedback while reaching to one of four targets in a center-out design. By the end of this block, participants were accurate in reaching toward the target (*t_19_* = 0.14, *p* = 0.89) with no systematic differences between targets (*F*_3,57_ = 0.29, *p* = 0.83; Fig. 2A,B). Following this Baseline block, rotations were introduced where each target location was associated with a unique angle of rotation (30°, 50°, 70°, and 90°) with the specific target-rotation pairing randomized across participants. Participants quickly began to adjust their reaches to offset the imposed rotations, with participants successfully adapting to all rotation magnitudes by the final epoch of the block (30°: *t_19_* = 0.11, *p* = 0.91; 50°: *t_19_* = −0.76, *p* = 0.46; 70°: *t_19_* = −1.48, *p* = 0.16; 90°: *t_19_* = −1.73, *p* = 0.10) with only marginal differences in performance errors between targets (*F*_3,57_ = 2.68, *p* = 0.055). Given the delay in endpoint feedback, this pattern is consistent with participants utilizing explicit strategies to adapt flexibly to multiple sets of visuomotor perturbations. After this Rotation block, visual feedback of the cursor and, consequently the imposed rotation, was removed, and participants were instructed to aim directly toward the target (No-Feedback “washout” block). Participants displayed a small, marginal aftereffect (3.7° ± 8.1°; *t_19_* = 2.06, *p* = 0.053; Fig. 2A) confirming that learning was the result of explicit strategies.

**Figure 2.**
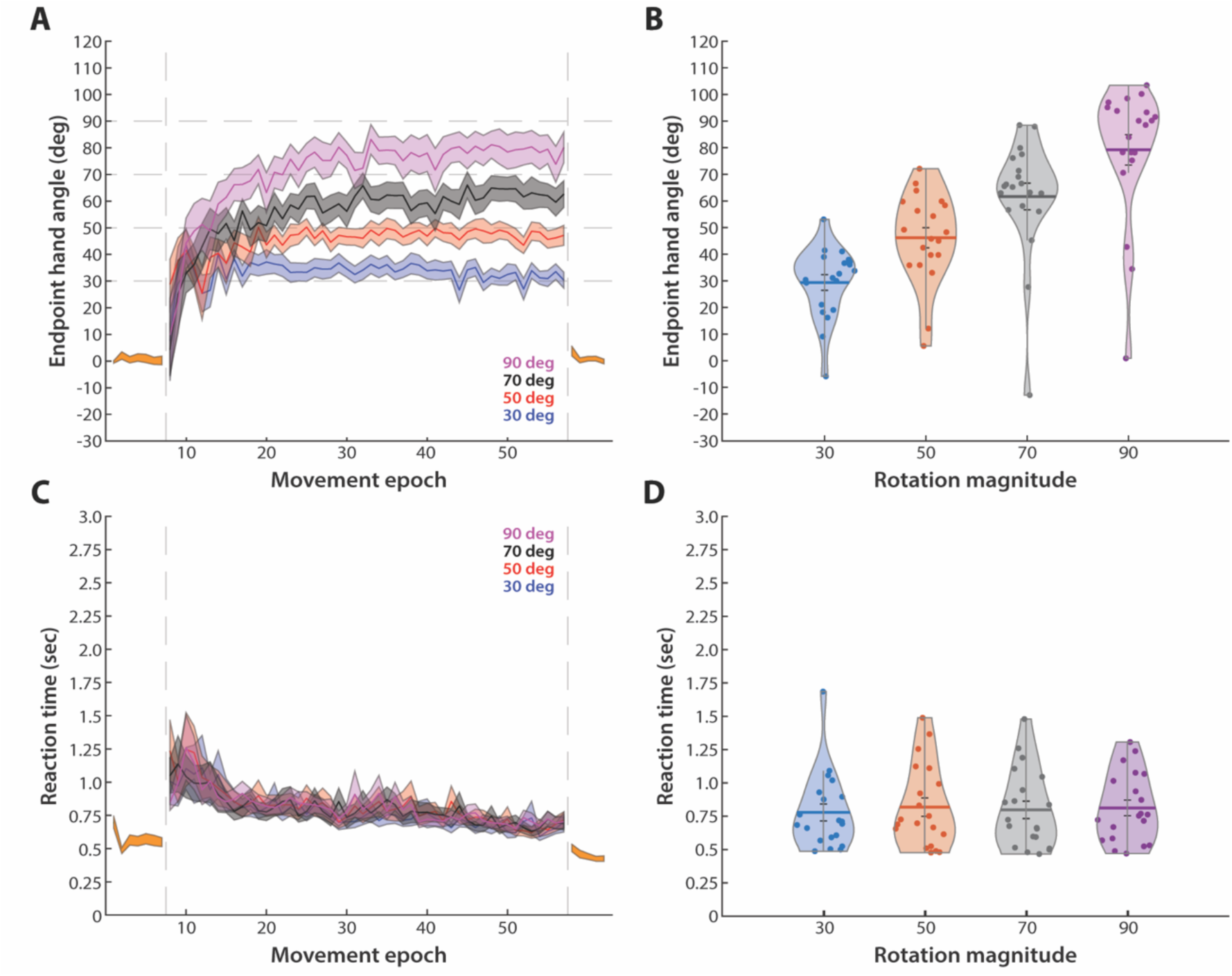
Behavior in experiment 1. (A, C) Endpoint hand angles (A) and RT (C) during the Baseline, Rotation and No-feedback blocks. Data during the Baseline and No-feedback blocks is shown in orange. Data during the Rotation block is separated by the magnitude of imposed rotation. Solid lines represent the mean, and the shaded ribbons represent SEM, across participants. Vertical dashed lines denote start and end of the Rotation block. (B) Endpoint hand angle in the final epoch of the Rotation block, separated by the magnitude of imposed rotation. (D) Mean RT across the Rotation block, separated by the magnitude of imposed rotation. Dots indicate individual participants, bars represent the mean across participants, and shaded violin regions show the distribution across the sample.

To determine if learning was the outcome of algorithmic- or retrieval-based strategies, we analyzed participants’ preparation time (reaction time; RT). If RTs scale with rotation magnitude, then it suggests that an algorithmic strategy was employed (McDougle & Taylor, 2019). Here, while RTs showed an initial sharp increase followed by a monotonic decrease over the course of the Rotation block, there were no differences in RTs as a function of rotation magnitude (*F*_3,57_ = 0.69, *p* = 0.56; Fig. 2C,D). This pattern is inconsistent with an algorithmic strategy and, instead, indicative of the use of an explicit memory retrieval strategy.

These findings suggest that learning to explicitly adapt to a complex set of sensorimotor mappings – more complex than has been previously studied with visuomotor adaptation tasks – poses little problem for participants. Because RTs did not scale with rotation magnitude, our findings suggest that participants learned to store and retrieve the solution appropriate to the relationship between target location and imposed rotation, from memory (McDougle & Taylor, 2019; Velazquez-Vargas & Taylor, 2024). This ability to learn and retrieve four distinct solutions is consistent with long-standing estimates of visual short-term working memory (VSTM) capacity, which is typically limited to three to four items (Cowan, 2001, 2010; Luck & Vogel, 1997; Zhang & Luck, 2008). As such, an open question is whether such capacity limits might represent a fundamental constraint on explicit strategies in complex settings. To answer this question, we sought to push well beyond such capacity limits by doubling the number of sensorimotor mappings. Specifically, in Experiment 2, participants were tasked with learning eight distinct target-rotation pairings. We should note that in a pilot study (Bejjanki et al., 2025), we found considerable individual differences across participants, when they were tasked with learning eight unique action-outcome mappings. As such, in order to adequately sample across-participant variability, we tripled the sample size to sixty participants.

### Experiment 2

The experimental task unfolded identical to Experiment 1, but with participants being tasked with attempting to counteract eight different rotations that were linked to eight unique target locations. By the end of the Baseline block, participants were accurate in reaching toward the targets (*t_59_* = −0.57, *p* = 0.57) with no systematic differences between targets (*F*_7,413_ = 0.38, *p* = 0.92; Fig. 3A). Following the Baseline block, rotations were introduced where each target location was associated with a unique angle of rotation (20°, 30°, 40°, 50°, 60°, 70°, 80°, and 90°) with the specific target-rotation pairing randomized across participants. In contrast to our findings in experiment 1, although they began to rapidly adjust their reaches to offset the imposed rotations, participants in experiment 2 failed to adapt to all the target-rotation pairs even by the last epoch of the Rotation block. Interestingly, this failure was not uniform: participants appeared to adapt to the 50° and 60° rotations appropriately – we found no difference between endpoint hand angle and the imposed rotation magnitude (both *p*s > 0.38); but for all other rotation magnitudes we found reliable (*p*s < 0.01 for 20°, 40°, 70°, 80°, and 90° rotations) or marginal differences (*p* = 0.08 for the 30° rotation).

**Figure 3.**
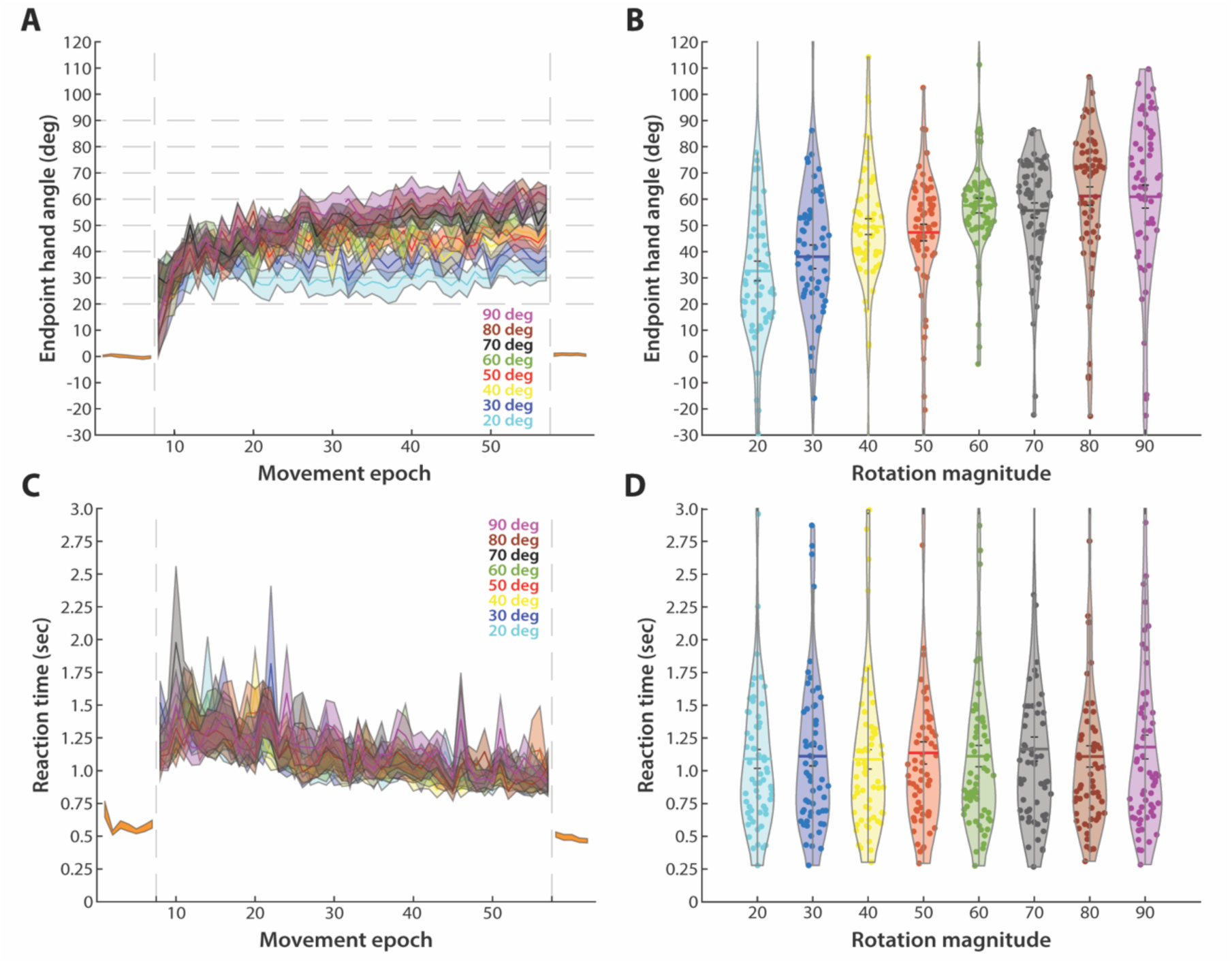
Behavior in experiment 2. (A, C) Endpoint hand angles (A) and RT (C) during the Baseline, Rotation and No-feedback blocks. Data during the Baseline and No-feedback blocks is shown in orange. Data during the Rotation block is separated by the magnitude of imposed rotation. Solid lines represent the mean, and the shaded ribbons represent SEM, across participants. Vertical dashed lines denote start and end of the Rotation block. (B) Endpoint hand angle in the final epoch of the Rotation block, separated by the magnitude of imposed rotation. (D) Mean RT across the Rotation block, separated by the magnitude of imposed rotation. Dots indicate individual participants, bars represent the mean across participants, and shaded violin regions show the distribution across the sample.

Indeed, the overall pattern is suggestive of participants tending to compensate for approximately the average of all the rotations, overshooting for rotations less than 50° and undershooting for rotations greater than 60°. At first glance, this pattern could appear to reflect a “compromise” solution where participants settled on an intermediate value strategy for all the target-rotation pairs. Indeed, the use of a similar solution across target-rotation pairs is supported by the lack of a significant relationship between rotation magnitudes and RTs (*F*_7,413_ = 1.99, *p* = 0.055; Fig. 3D), which could be the result of leveraging the same algorithmic or retrieval strategy for each target-rotation pair. However, this possibility would predict a flat relationship between rotation magnitude and endpoint hand angle; instead, as can be seen in Figure 3B, there is a clear slope across rotation magnitudes (*F*_7,413_ = 12.26, *p* <0.001). This raises the question as to why participants’ behavior would settle on an approximate intermediate solution, that is modulated in a manner that is specific to each rotation magnitude. We address this question in the next section by inspecting individual differences.

Finally, after the Rotation block, the rotation and cursor feedback were removed, and participants were instructed to aim directly toward the target (No-Feedback block). As in experiment 1, participants once again displayed a very small (0.6 ± 4.6 degrees) aftereffect which was not reliably different from zero (*t_59_* = 1.06, *p* = 0.29; Fig. 3A). This confirms that the limited adaptation we observed here is the result of explicit re-aiming strategies and not implicit recalibration.

### Individual differences in learning

The group-averaged performance in Experiment 2 (with eight unique target-rotation pairs), suggests that when pushed beyond working memory capacity, participants converged toward the mean of the imposed perturbations. However, this apparent “averaging” may not reflect the behavior of any single individual. One possibility is that different participants successfully adapted to different subsets of target-rotation pairs, such that when combined across individuals, the aggregate behavior appeared centered around the mean rotation value. To evaluate this possibility, and to more precisely characterize individual learning curves, we fit each participant’s learning function for each target-rotation pair with a three-parameter Weibull function (see Methods). According to Gallistel et al. (2004), the Weibull function provides a quantitative way to capture the latency, rate, and asymptote of individual learning functions, which avoids potential distortions that can arise when asynchronous or idiosyncratic learning functions are averaged across participants.

The fits captured the learning dynamics modestly well across participants (*r* = 0.59±0.06) despite considerable variability across participants (Fig. 4). Consistent with this variability, we found a wide range of asymptotic performance levels: some participants achieved near-complete compensation across all eight rotations (Fig. 4A), others plateaued at intermediate levels consistent with partial adaptation (Fig. 4B), and a subset showed minimal learning despite continued engagement throughout the task (i.e., they did not simply quit trying; Fig. 4C). Importantly, it does not appear that different participants learned different subsets of target-rotation pairs (Fig. 4D). Furthermore, the asymptotic performance values mirror the trend observed with the endpoint hand angles at the end of the Rotation block: performance scales with rotation magnitude but overshoots for smaller rotations and undershoots for larger rotations for the vast majority of participants (Fig. 4E). Furthermore, these performance gains are obtained relatively early in training, with an average latency of 1-2 epochs into the Rotation block (Fig. 4F), and with slopes that are all much greater than 1, which is more consistent with a sigmoidal function (i.e., “aha” moments) as opposed to a more gradual process. Taken together, these results suggest that the group-averaged “averaging” pattern might reflect the aggregate of individual differences in learning capacity rather than a top-down averaging strategy *per se*. It remains an open question whether these performance differences across participants is the result of individual differences in working memory (Christou et al., 2016).

**Figure 4.**
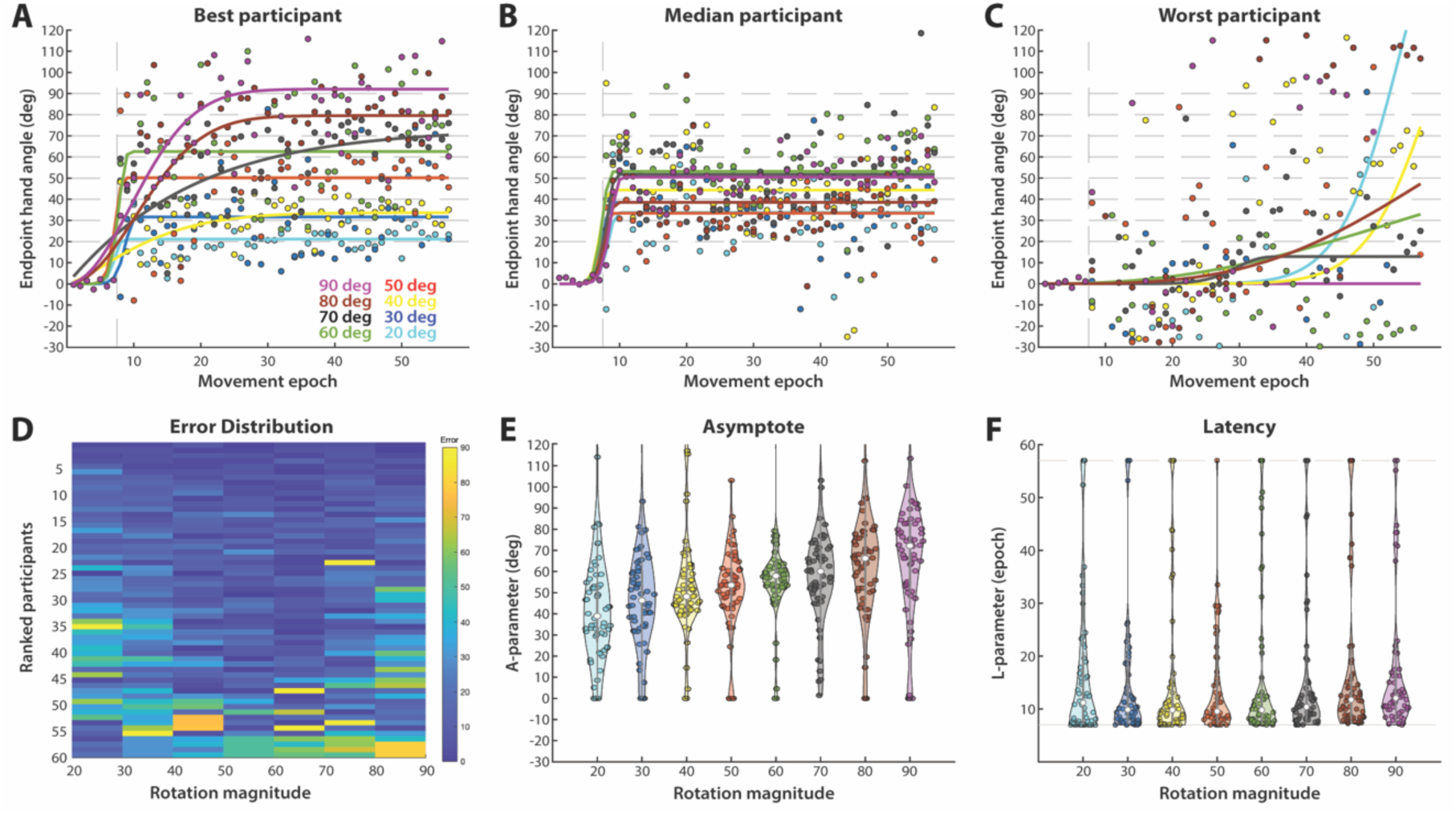
Individual differences in experiment 2. (A-C) Endpoint hand angles during the Baseline and Rotation blocks for the best (A), median (B) and worst participant (C) in our sample. Dots indicate individual trials, and solid lines represent best-fit Weibull functions. Colors depict imposed rotation magnitude. (D) Distribution of errors in endpoint hand angles in the final epoch of the Rotation Block. Colors depict error (see color bar), separated by rotation magnitude (x-axis) and participants (y-axis; participants ranked by overall error from low [top] to high [bottom]). (E) Distributions of the asymptote parameter A and (F) latency to learn parameter L for best-fit Weibull functions across participants and rotation magnitudes (colors same as in A-C). Dots indicate individual participants, and shaded violin regions show the distribution across the sample.

### Model Simulations

The variability in asymptotic performance across participants in Experiment 2, which scales with rotation magnitude but trends toward the mean of the imposed rotations, suggests that learners may not be able to maintain all eight target–rotation pairs associations concurrently. Instead, performance may reflect the limits of a finite (spatial) working memory buffer that can only hold a subset of currently relevant mappings. This idea is consistent with current accounts of working memory capacity, which estimate an upper bound of roughly 4 ± 1 items (Cowan, 2001). If explicit re-aiming strategies rely on such a limited capacity, then as the number of required mappings exceeds this capacity, participants should begin to lose access to specific stimulus-response (i.e., target-rotation pair) associations that are outside their working memory “window”, and result in guessing (Zhang & Luck, 2008). Inspired by this idea, we developed a simple, parameter-free working memory model which assumes that, at any given time, a participant maintains a small, fixed number of target–rotation associations in an active working memory window (Fig. 5A). Thus, in a given trial, if the mapping relevant to the target is available within this window, the appropriate strategy is implemented. However, if the target falls outside this limited window, the participant is forced to guess, which is modeled as a random draw from all the mappings that are not currently available in the working memory store. We used this model to simulate behavior for the final epoch of eight trials, for each of 60 model “participants” (i.e., matched to the sample size used in Experiment 2), thereby allowing us to compare model output to the behavior observed in Experiment 2 at the end of the rotation block. Finally, we varied the size of the working memory window (from one “slot” to eight “slots”), in order to elucidate the influence of varying capacity limits on behavior.

**Figure 5.**
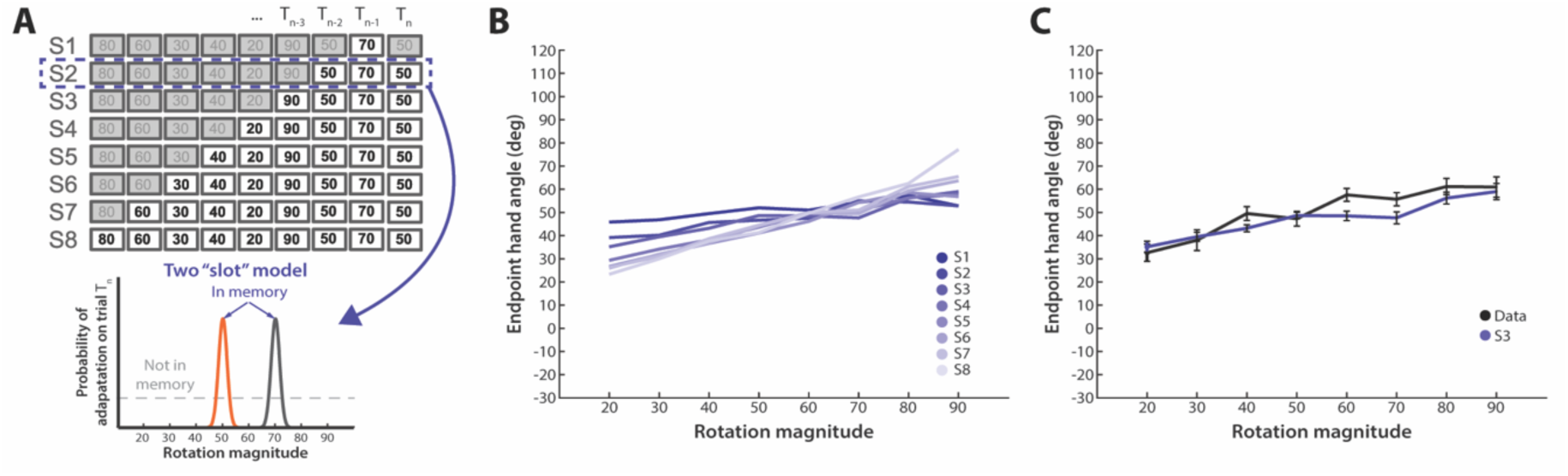
Model of working memory. (A) On a given trial, a model “participant” maintains a small, fixed number of target–rotation associations in an active working memory window. *Top:* Illustration showing solutions cached in working memory for models with one “slot” [S1] to eight “slots” [S8]. *Bottom:* Illustration showing probability of participant successfully adapting to the imposed rotation on a given trial, based on the active working memory cache, for the two “slot” model [S2]. If the mapping relevant to the target is available in the cache, the appropriate strategy is implemented. If the target falls outside this limited window, the participant is forced to guess, which is modeled as a random draw from all the mappings that are not currently available in the working memory store. (B) Simulated behavior in the final epoch of the Rotation block, averaged across 60 model “participants”, for models S1 to S8. (C) Behavior of participants in our sample for experiment 2 is best predicted by a model with three “slots” [S3]. Solid lines represent the mean, and error bars represent SEM, across participants.

When the model only included one slot in working memory, we found that the group-averaged behavior collapsed approximately to a mean rotation value, as would be expected given that “participants” are primarily guessing - i.e., drawing randomly from the available mappings. However, as the number of available working memory slots increased, the group-averaged behavior increasingly demonstrated a slope across rotation magnitudes, with a consistent overshooting for small rotations and undershooting for large rotations. Once again, the inclusion of random guesses for mappings not in the working memory store provides a clear explanation for this pattern (Fig. 5B). Finally, we found that the limited behavior observed in Experiment 2, wherein participants were unable to adapt to all eight target-rotation pairs, is best predicted (minimal mean RMS error) by this model when it includes three slots (Fig. 5C). It is worth noting that at no point (i.e., even with the maximum eight slots) did our model output reflect perfect performance - i.e., where the model output perfectly matched the imposed rotation angle. This is a consequence of our assumption that multiple slots within the available working memory store could be occupied by the same target-rotation mapping. This meant that even with eight slots, model “participants” were still occasionally forced to guess, which depressed the slope away from the unity line.

Taken together, these model simulations provide a mechanistic explanation for the bias towards the average rotation value observed in experiment 2 – i.e., that this pattern might be best explained as a product of a combination of limited working memory capacity and random guessing.

## Discussion

The present work set out to test the extent to which explicit strategies can support visuomotor adaptation in complex settings. While the traditional view of sensorimotor adaptation has long emphasized cerebellar-dependent implicit recalibration (Miall & Wolpert, 1996; Shadmehr et al., 2010), converging evidence indicates that much of the performance improvements observed in visuomotor adaptation tasks arise from deliberate, explicit strategies. However, while explicit re-aiming strategies can produce rapid and flexible improvements in task performance (Bond & Taylor, 2015; Taylor et al., 2014), they also rely on cognitive resources, such as working memory and executive control, that are inherently capacity-limited (Anguera et al., 2010; McDougle & Taylor, 2019; Taylor & Ivry, 2014; Velazquez-Vargas & Taylor, 2024). Thus, here we asked whether such re-aiming strategies could handle complex sensorimotor mappings that are at least a step closer to those found in real-world settings, where multiple distinct action–outcome relationships must be learned and maintained simultaneously. We found that participants could successfully learn and retrieve up to four distinct visuomotor mappings, which is consistent with classic estimates of visual working-memory capacity (Cowan, 2010; Zhang & Luck, 2008); however, when task complexity doubled to eight mappings, participants failed to adapt to all the target-rotation pairs. This failure was systematic – participants tended to compensate for approximately the average of the rotation magnitudes, overshooting for rotations less than 50° and undershooting for rotations greater than 60°.

### Influence of working memory constraints

The particular form of working memory limitation we observed, which resulted neither in total failure nor idiosyncratic learning, but instead in apparent averaging across perturbation magnitudes, provides insight into the underlying limitation. Aggregating across trials yields an average behavior that surprisingly scales proportionally with the magnitude of the perturbation but tends toward the average of the rotation set. This results in a systematic overshoot–undershoot slope that could be viewed as a solution that compromises error loss. However, it does not appear to be the result of visuomotor mental rotation, as participants’ RT did not proportionally scale with rotation magnitude. Instead, the scaling appears to naturally arise from what would be predicted if behavior is subject to a combination of limited working memory capacity and random guessing. Indeed, in support of this notion, participants’ behavior in Experiment 2 is consistent with the output from our parameter-free working-memory model. The model assumed a small trial-by-trial moving window of active representations (3 items). When a target’s mapping fell within this window, the correct compensatory response was implemented, but when it fell outside, the response was effectively guessed. This is consistent with our previous work that found that participants could hold only a couple of target-rotation strategies in working memory (Velazquez-Vargas & Taylor, 2024).

Recently the role of working memory in sensorimotor adaptation has gained much attention (Ebrahimi & Ostry, 2024; McDougle & Hillman, 2025; Sidarta et al., 2018). While much of this work has focused on somatosensory memory (Ebrahimi & Ostry, 2024; Sidarta et al., 2018), the pattern we observe here is not easily explained by somatosensory working-memory: the imposed perturbation provided no proprioceptive signal, visual error information was decoupled from participants’ movements, and the time between successive trials at specific target locations was quite long. Instead, our findings are consistent with the notion that the limitation is likely the result of visual short-term or spatial working memory. Or, more intriguingly, these effects may reflect a domain-specific “motor working memory” – a distinct subsystem proposed to maintain abstract, action representations (Hillman et al., 2024; McDougle & Hillman, 2025). Indeed, prior research has shown that individual differences in explicit learning during visuomotor adaptation correlate with estimates of effector-independent motor working memory capacity (Hillman et al., 2025).

An open question concerns whether such capacity limits can be overcome with extensive practice. One possibility is that repeated exposure allows explicit associations to be consolidated into long-term memory, effectively expanding usable capacity over time. Indeed, in previous work we found that if the target-rotation pairs were held consistent – even when exceeding working memory capacity – participant’s accuracy improved (Velazquez-Vargas & Taylor, 2024). However, this previous study was at the lower edge of working memory capacity (5 items), and thus may have presented participants with enough repetition to enable transfer to long-term memory. In our current experiment, when we far exceeded this limit, we found that, at least over the timescale of a typical adaptation experiment, participants’ performance seemed to asymptote.

Our results therefore suggest there may be a potential “sweet spot” or desirable level of difficulty for learning (Al-Fawakhiri et al., 2023; Bjork & Bjork, 2011; Wilson et al., 2019), especially when the task demands substantial contributions from higher-order cognition. When the task complexity lies within working-memory capacity, participants can flexibly retrieve and update mappings, producing rapid learning. On the other hand, when complexity exceeds capacity, learning struggles to take hold and performance asymptotes. This could explain why difficult motor learning tasks are often decomposed into components or subroutines, otherwise they may overwhelm available resources and blunt learning (Correa et al., 2023; Griessbach & Haith, 2025; Velázquez-Vargas & Taylor, 2025).

### Alternative avenues for learning

If explicit strategies can quickly lose their utility when task complexity exceeds working memory capacity limitations, are there other learning systems that could step into the breach? Reinforcement learning (RL) is an obvious candidate, which could, in principle, support incremental learning of stimulus–response mappings through trial-and-error updating of value functions. Yet RL mechanisms, at least in situations relevant to the current task, operate very slowly. Indeed, in simple associative learning tasks where the stimulus-response set size exceeds working memory capacity, although learning via RL is possible, it is very slow (Collins & Frank, 2012, 2018). This could explain why motor learning for highly complex skills often takes years to master.

A form of statistical inference offers another possibility. Humans can adaptively adjust learning rates based on environmental volatility, thereby efficiently integrating prior expectations with new evidence (Behrens et al., 2007; Nassar et al., 2012; Nassar et al., 2010; Razmi & Nassar, 2022). However, in the present context, the underlying generative function linking the different target-rotation pairs is arbitrary, providing no discoverable relationship between action-outcome mappings. In contrast, if the set of perturbations followed a lawful transformation (e.g., a smooth function over target angle), an algorithmic process could, in principle, approximate that function through interpolation or regression-like operations. Indeed, in earlier work, we found that the motor system could adapt an internal model when the spatial frequency of perturbation functions was relatively smooth (Thoroughman & Taylor, 2005). In addition, explicit strategies could scale more effectively if the underlying function were even minimally stationary, allowing for structure learning over the class of perturbations (Bond & Taylor, 2017; Braun et al., 2009; Braun et al., 2010). Such learning would likely not rely on memory-based retrieval, as we have found here, but instead rely on algorithmic strategies, such as mentally rotating a planned movement vector. Future work will be necessary to determine how task complexity might limit the use of algorithmic strategies for visuomotor adaptation.

### Complex sensorimotor mappings and learning de novo

An important conceptual implication of the present results concerns the distinction between adaptation and *de novo* learning (Krakauer et al., 2019). Adaptation is typically defined as the incremental recalibration of an existing control policy driven by sensory prediction errors, which is an automatic process that operates continuously and largely outside awareness. In contrast, *de novo* learning involves constructing a novel control policy, often requiring explicit hypothesis testing, flexible action selection, and the formation of new stimulus–response associations (Hadjiosif et al., 2021; Haith et al., 2015; Haith et al., 2022; Morehead et al., 2017; Yang et al., 2021). The current paradigm, which requires storing and retrieving multiple arbitrary re-aiming rules, fits more squarely under *de novo* learning of a sensorimotor mapping.

Consider for example, mirror-reversal learning. When faced with a mirrored mapping, implicit adaptation proceeds in the *wrong* direction, reflecting an inappropriate attempt to recalibrate an existing forward model rather than construct a new one (Hadjiosif et al., 2021; Lillicrap et al., 2013; Telgen et al., 2014; Wilterson & Taylor, 2021). Successful performance therefore depends on developing a new, explicitly specified mapping that is gradually proceduralized with practice (Huberdeau et al., 2015). The qualitative features of this process – slow acquisition, abrupt transitions, heavy reliance on working memory, and the formation of discrete rules – closely resemble the learning patterns observed in the present task when the number of distinct rotations increases beyond a small set.

Additional support for this view comes from tasks requiring the acquisition of arbitrary visuomotor associations or novel coordination policies. In tasks involving spatially distributed cue–action mappings, learning unfolds slowly and often in a fragmented manner, with step-like improvements suggesting the acquisition of discrete action rules rather than smooth, error-based updating (Bapi et al., 2000; Bera et al., 2021; Collins & Frank, 2013; Fermin et al., 2010; Velazquez-Vargas & Taylor, 2024; Velázquez-Vargas & Taylor, 2025). Similarly, in tasks requiring the formation of new bimanual coordination patterns, performance is only successful when the mapping complexity remains within a limited capacity – approximately four independent “chunks” – and deteriorates sharply once this limit is exceeded (Yokoi et al., 2018). These hallmarks echo the constraints seen here: participants can acquire a set of a few independent re-aiming mappings, but performance collapses once the number of required mappings exceeds working-memory capacity.

Taken together, these convergent findings suggest that the limitations we observe do not reflect properties of adaptation but instead reflect general constraints on *de novo* skill learning under heavy cognitive load. Adaptation mechanisms do not exhibit such hard capacity limits and tend to fail gradually rather than discretely. In contrast, constructing a new visuomotor mapping requires maintaining and retrieving explicit action rules, a process bounded by domain-general working-memory and executive-control systems. This perspective clarifies why “adaptation” measured under conditions dominated by explicit re-aiming is better characterized as a form of *de novo* learning: its success depends on rule formation and memory-based retrieval rather than implicit recalibration.

## Conclusion

By identifying a cognitive bandwidth limit on the utility of explicit re-aiming strategies for supporting visuomotor adaptation in complex settings, this work refines our understanding of how different processes can usefully support learning under heavy cognitive load. Explicit strategies are effective for improving performance early on, at least when task complexity is within the capacity of working memory. However, as our results demonstrate, their utility sharply declines once task complexity exceeds working-memory capacity. Thus, the same cognitive resources that confer flexibility also impose a hard constraint on scalability. This observation has implications for how we think about the real-world usefulness of explicit strategies. Everyday motor behaviors – from tool use to musical performance – often require maintaining and switching among many distinct mappings. In such cases, our results suggest that reliance on explicit control alone would be computationally infeasible. Instead, when moving beyond even a modest level of complexity, learners must recruit slower, more automatic learning processes, or restructure the task so that it becomes compositional or chunked into manageable subroutines.

## Methods

### Experiment 1

#### Participants

20 undergraduates [9 men, 11 women; mean age 19.9 years (SD 1.33)] participated in this experiment. Participants received course credit. All participants had normal or corrected-to-normal vision and were right-handed. The protocol was approved by the IRB at Princeton University, and all participants provided written informed consent.

#### Experimental Apparatus

Visual stimuli were displayed on a horizontally mounted 21.5-in. touch-sensitive monitor (60 Hz refresh rate; Fig. 1A). Participants’ reaches were sampled at 60 Hz by a digitizing pen as they slid their hand across the surface of a tablet mounted 25 cm below the monitor. The game was controlled by a Dell desktop PC running custom MATLAB and Psychtoolbox software (Brainard, 1997).

#### Procedure

Participants began each trial with their right hand at the center of the visual workspace (Fig. 1A). To aid them in finding the center, a white circle expanded or contracted with the radial distance of their hand position from the center of the tablet. Once their hand was 10mm from the center of the start area (0.3cm radius), a white circular cursor (0.15cm radius) appeared. After holding this position for 300ms, a circular green target (0.25cm radius) appeared 7cm from the start position. Participants were instructed to make a reaching movement by “shooting” their right hand through the target. If the reach duration exceeded 600ms, participants received a “too slow” auditory warning. Only end-point feedback, presented for 500ms as a cursor position on the virtual ring (7 cm), was provided for each reach (except for the first 8 trials of the Baseline Block, which provided continuous online feedback). Specifically, for all remaining trials in the Baseline Block, the white cursor was removed when the participant’s hand exceeded a radial distance of 0.4 cm from the start position and reappeared when their hand passed the virtual ring, along with a “thunk” sound that let them know that they had reached far enough. In order to limit implicit recalibration and better isolate explicit learning, endpoint feedback was delayed on each trial by one second (Brudner et al., 2016; Kitazawa et al., 1995; Schween & Hegele, 2017).

All conditions followed the same five-block structure. Participants reached to one of four targets (located at 15°, 105°, 195° or 285°), each randomly associated with a different visuomotor rotation magnitude (±30°, ±50°, ±70° or ±90°). For the first 8 trials of the Baseline Block, participants were provided with veridical, continuous online feedback to familiarize them with the task. For the next 8 trials, online feedback was replaced by endpoint feedback. For the remainder of the Baseline Block (40 trials), participants were again provided only with endpoint feedback but this feedback was delayed by one second. A clockwise/counter clockwise rotation (counterbalanced across participants) was introduced in the fourth block of 400 trials (Rotation block). In the fifth block of 40 trials, visual feedback of the cursor was removed (No-Feedback “washout” block), and participants were provided with specific aiming instructions: “Please aim directly to the green target. You won’t see any cursor feedback. You’ll hear a “thunk” sound that lets you know that you have reached far enough.”

#### Data Analysis

Kinematic and statistical analyses were performed with MATLAB (MathWorks, Natick, MA). To assess performance, we examined participants’ endpoint hand angle measured when movements passed a radial distance of 7 cm. Each movement trajectory was transformed from Cartesian to polar coordinates and rotated to a common reference axis with the target location set at 0°. Trials were separated by rotation during the rotation block, and endpoint hand angles for each rotation were averaged in eight-trial bins (epochs) for further analyses, with each epoch including an average across two trials. We opted to consider epochs spanning eight trials because our study included up to eight target locations, which were pseudo-randomized. Moreover, since the sequence of target locations was randomized across participants, this procedure removes biases associated with specific target locations.

We examined participants’ reach locations in the last epoch in the rotation block for each rotation. We submitted the difference between the endpoint hand angles in the last epoch and the magnitude of the rotation, to evaluate the extent of learning. Note that in our design, participants’ endpoint hand angle directly represented their explicit aim angle. To quantify any aftereffect, we submitted the first epoch of the No-Feedback block to a one-sample *t*-test. Note, to remove potential biases in reaching, we subtracted the endpoint hand angles from the final epoch in the Baseline block, from all subsequent phases, as is custom in visuomotor rotation studies. Finally, for each participant and target-rotation pair, we computed the average reaction time (RT) across the entire Rotation block. RT was measured as the time elapsed from target appearance to the hand leaving the start area. We submitted these measures to a one-way repeated measures ANOVA to evaluate the extent to which participants’ RTs varied as a function of the rotation magnitude.

### Experiment 2

#### Participants

60 undergraduates [28 men, 29 women, 3 other/unknown; mean age 19.67 years (SD 1.20)] participated in this experiment. Participants received course credit. All participants had normal or corrected-to-normal vision and were right-handed. The protocol was approved by the IRB at Princeton University, and all participants provided written informed consent.

#### Procedure, Experimental Apparatus, and Analysis

The task procedure, experimental apparatus, and analyses were nearly identical to that used in Experiment 1 except for the fact that participants now reached towards one of eight targets (located at 15°, 60°, 105°, 150°, 195°, 240°, 285° or 330°), each randomly associated with a different visuomotor rotation magnitude (±20°, ±30°, ±40°, ±50°, ±60°, ±70°, ±80°, or ±90°).

#### Data Analysis

Analyses were similar to Experiment 1 with the exception that we also fit the Weibull function fits to individual learning curves (Gallistel et al., 2004):

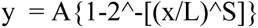

where an individual participant’s hand angle (y) is captured by an asymptotic learning parameter (A), latency of when learning begins (L), and steepness (S) of the learning function over the course of training (x).

Prior to fitting each participant, we performed two sensitivity analyses: 1) to determine if baseline trials should be included in the fitted time series, and 2) to set an upper bound on the unbounded steepness parameter. This first analysis revealed that the inclusion of all baseline trials minimized root-mean-square (RMS) error, presumably because it provides a clearer picture of change points (or “aha” moments). The second analysis revealed that 95% of the asymptotic improvement in RMS error was achieved with an upper bound of S = 6.26. The greater the S-value, the steeper the function and for most of the participants the best fitting S parameter was at the limit, which supports prior work suggesting that explicit adaptation is abrupt (Taylor & Ivry 2012; Taylor et al., 2014; Tsay, Kim, McDougle, Taylor et al., 2024; Townsend et al., 2025). Following these sensitivity analyses, each participant’s learning curve was fit with the bounded Weibull function using the *fmincon* function in MATLAB.

## Acknowledgements

We wish to thank Chandra Greenberg for assistance with data collection, and Deanna Durben and Kelly Grossman for help with preliminary experiments that informed this study. This work was supported by a Faculty Teacher-Scholar Award from Hamilton College to Vikranth R. Bejjanki and grant R01NS131552 from the National Institute of Neurological Disorders and Stroke (NINDS) of the National Institutes of Health (NIH) awarded to Jordan A. Taylor. The funders had no role in study design, data collection and analysis, decision to publish, or preparation of the findings.

